# Loss of endothelial glucocorticoid receptor accelerates organ fibrosis in *db/db* mice

**DOI:** 10.1101/2023.03.20.533532

**Authors:** Swayam Prakash Srivastava, Julie E. Goodwin

## Abstract

Endothelial cells play a key role in maintaining homeostasis and are deranged in many disease processes, including fibrotic conditions. Absence of the endothelial glucocorticoid receptor (GR) has been shown to accelerate diabetic kidney fibrosis in part through up regulation of Wnt signaling. The db/db mouse model is a model of spontaneous type 2 diabetes that has been noted to develop fibrosis in multiple organs over time, including the kidneys. This study aimed to determine the effect of loss of endothelial GR on organ fibrosis in the *db/db* model. Db/Db mice lacking endothelial GR showed more severe fibrosis in multiple organs compared to endothelial GR-replete db/db mice. Organ fibrosis could be substantially improved either through administration of a Wnt inhibitor or metformin. IL-6 is a key cytokine driving the fibrosis phenotype and is mechanistically linked to Wnt signaling. The db/db model is an important tool to study mechanisms of fibrosis and its phenotype in the absence of endothelial GR highlights the synergistic effects of Wnt signaling and inflammation in the pathogenesis or organ fibrosis.

## Introduction

Despite the rising incidence of fibrotic kidney disease, driven to a great extent by a continued increase in the prevalence of diabetes (1, 2), molecular mechanisms of renal fibrosis remain incompletely understood. One important tool to study this phenomenon is the C57BL/6 *db/db* mouse model which demonstrates type 2 diabetes, significant albuminuria, renal histological damage and reduced renal function by 32 weeks of age (3). Studies of *db/db* mice have demonstrated that these animals display deranged composition and morphology of key organs such as kidney, liver, heart and adipose tissue as well as significantly different viscoelastic properties, even when compared to the *ob/ob* mouse, which also possesses a leptin mutation (4), highlighting the fact that biomechanical properties may be involved in the pathogenesis of diabetic complications. Additional insights gained through the study of the *db/db* model have revealed other mechanistic underpinnings of diabetes including oxidative stress, advanced glycation end products, renin-angiotensin-aldosterone (RAAS) activation and unchecked inflammation (3).

Previously, we demonstrated that mice lacking the endothelial glucocorticoid receptor (*GR^ECKO^*) were highly susceptible to streptozotocin-induced diabetic nephropathy and demonstrated augmented Wnt signaling, aberrant cytokine reprogramming and suppressed fatty acid oxidation (FAO) (5, 6), leading to the conclusion that the endothelial glucocorticoid receptor (GR) is a key regulatory molecule in the pathophysiology of renal fibrosis. Evolving understanding of the complicated Wnt signaling pathway suggests a favorable role in kidney repair when transiently activated, but a detrimental effect which promotes injury and fibrosis in states of unremitting Wnt activation (7). In this work, we show that when bred onto the *db/db* background, *GR^ECKO^* mice display widespread organ fibrosis; high-fat diet (HFD) feeding of *GR^ECKO^* mice can produce a similar phenotype. This phenotype can be partially rescued by either administration of metformin or a small molecule Wnt inhibitor. Further, IL-6 is a key cytokine driving fibrosis in this model. We conclude that both the hyperinflammatory state and the concurrent uncontrolled augmented Wnt signaling that result from loss of endothelial GR (6, 8, 9) are further exacerbated by either HFD feeding or the *db/db* background. The *db/db*; *GR^ECKO^* model is a highly unique model which illustrates the key importance of both inflammation and Wnt signaling in the pathophysiology of diabetic nephropathy.

## Methods

### Animal experimentation

All experiments were performed according to a protocol approved by the Institutional Animal Care and Use Committee at the Yale University School of Medicine and were in accordance with the National Institute of Health (NIH) Guidelines for the Care of Laboratory Animals. Mice were housed at an ambient temperature of 68–79 °F with a humidity that ranged between 30 and 70%. They were exposed to 12-h light–dark cycles. Mice lacking endothelial GR (*GR^ECKO^*) were successively bred to heterozygous *Lepr^+/db^* mice to achieve the target mice: *db/db*; *GR^ECKO^*. Wnt inhibitor (LGK974) was provided to 16-week old male mice by gavage using a dose of 5 mg/kg at a frequency of six doses per week for 8 weeks, as described previously (10). Metformin was provided at 100 mg/kg by gavage for 8 weeks to some mice. In other experiments mice were received a HFD containing 40% fat (Research Diets, D12108). For glucose tolerance testing (GTT), mice were fasted overnight. On the following day, blood glucose profiles were measured at 0 min (baseline) and then at 30, 60, 90 and 120 min post oral glucose load of 3 g/kg body weight. Aside from this overnight fast for GTT, all mice had free access to food and water during experiments. Blood was obtained by retro-orbital bleed during experiments. Blood glucose was measured by glucose-strips. Tissues and blood were harvested at the time of sacrifice. Some tissues were minced and stored at −80 °C for gene expression and protein analysis. Other tissues were placed immediately in optimal cutting temperature compound for frozen sections or 4% paraformaldehyde for histologic staining.

### Morphological evaluation

Masson’s Trichrome-stained images were evaluated by ImageJ software, and the fibrotic areas were estimated. Deparaffinized sections were incubated with picrosirius red solution for 1 h at room temperature. The slides were washed twice with acetic acid solution for 30 seconds per wash. The slides were then dehydrated in absolute alcohol three times, cleared in xylene, and mounted with a synthetic resin. Sirius red staining was analyzed using ImageJ software, and fibrotic areas were quantified.

### In vitro experiments and siRNA transfection

HUVECs were used at passages four to eight and cultured in Endothelial Basal Medium-2 media with growth factors and 10% serum. Human GR-specific siRNA (Invitrogen) was used at a concentration of 100 nM for 48 h to effectively knockdown GR. Cells were treated with 10 μg/ml IgG control or IgG N-IL-6 (Bio X Cell), Wnt3a 200 ng/ml (Peprotech) or DKK-10 μg/ml (R&D systems) according to the experimental plan. When the cells reached 70% confluence, conditioned media from control siRNA- and GR siRNA-transfected HUVECs was tested for IL-6 expression.

### RNA isolation and qPCR

Total RNA was isolated using standard Trizol protocol. RNA was reverse transcribed using the iScript cDNA Synthesis kit (Bio-Rad) and qPCR was performed on a Bio-Rad C1000 Touch thermal cycler using the resultant cDNA, as well as qPCR Master mix and gene-specific primers. Results were quantified using the delta–delta-cycle threshold (Ct) method (ΔΔCt). All experiments were performed in triplicate and 18S was utilized as an internal control. The following primers were used:

α-SMA: Forward 5’-CTGACAGAGGCACCACTGAA and Reverse 5’-GAAATAGCCAAGCTCAG
FSP-1: Forward 5’-TTCCAGAAGGTGATGAG and Reverse 5’-TCATGGCAATGCAGGACAGGAAGA
Axin2: Forward 5’ AACCTATGCCCGTTTCCTCTA and Reverse 5’-GAGTGTAAAGACTTGGTCCACC
18S: Forward 5’-TTCCGATAACGAACGAGACTCT and Reverse 5’-GGCTGAACGCCACTTGTC

All primers were synthesized by the Keck Oligo Synthesis facility at Yale School of Medicine.

### Cytokine measurements

IL-1β, IL-6 and IFN-γ were measured by cytokine array using the Luminex platform.

### Statistical significance

All values are expressed as means ± SEM and analyzed using the statistical package for GraphPad Prism 7 (GraphPad Software, Inc., La Jolla, CA). One-way ANOVA, followed by Tukey’s test and two-way ANOVA was employed to analyze significance when comparing multiple independent groups. In each experiment, n represents the number of separate experiments (in vitro) or the number of mice (in vivo). Technical replicates were used to ensure the reliability of single values. The data were considered statistically significant at p < 0.05.

## Results

### Loss of endothelial GR causes organ fibrosis in db/db mice

We bred GR^ECKO^ mice onto the *db/db* background and assessed post-prandial glucose measurements after a 12-hour fast over a 2-week period. Control *db/db* mice and *db/db*; *GR^ECKO^* mice demonstrated similar serum glucose levels over a 14-day period (Figure 1A) as well as similar values during an oral glucose tolerance test after a glucose load of 3 mg/g body weight (Figure 1B,C). Further phenotyping showed that control *db/db* and *db/db*; *GR^ECKO^* mice had similar body weight, epididymal fat weight and liver weight, but *db/db*; *GR^ECKO^* mice had higher kidney weight/body weight ratios and increased heart weight (Figure 1D). Histologic examination of the kidneys and hearts from these animals clearly showed more fibrosis (MTS) and collagen deposition (Sirius Red) in organs from the *db/db*; *GR^ECKO^* mice (Figure 1E,F). Interestingly liver and adipose tissue from *db/db*; *GR^ECKO^* mice also showed more fibrosis compared to *db/db* controls, even though there was no change detected in these organs’ weight (Figure 1G,H).

**Figure 1:**
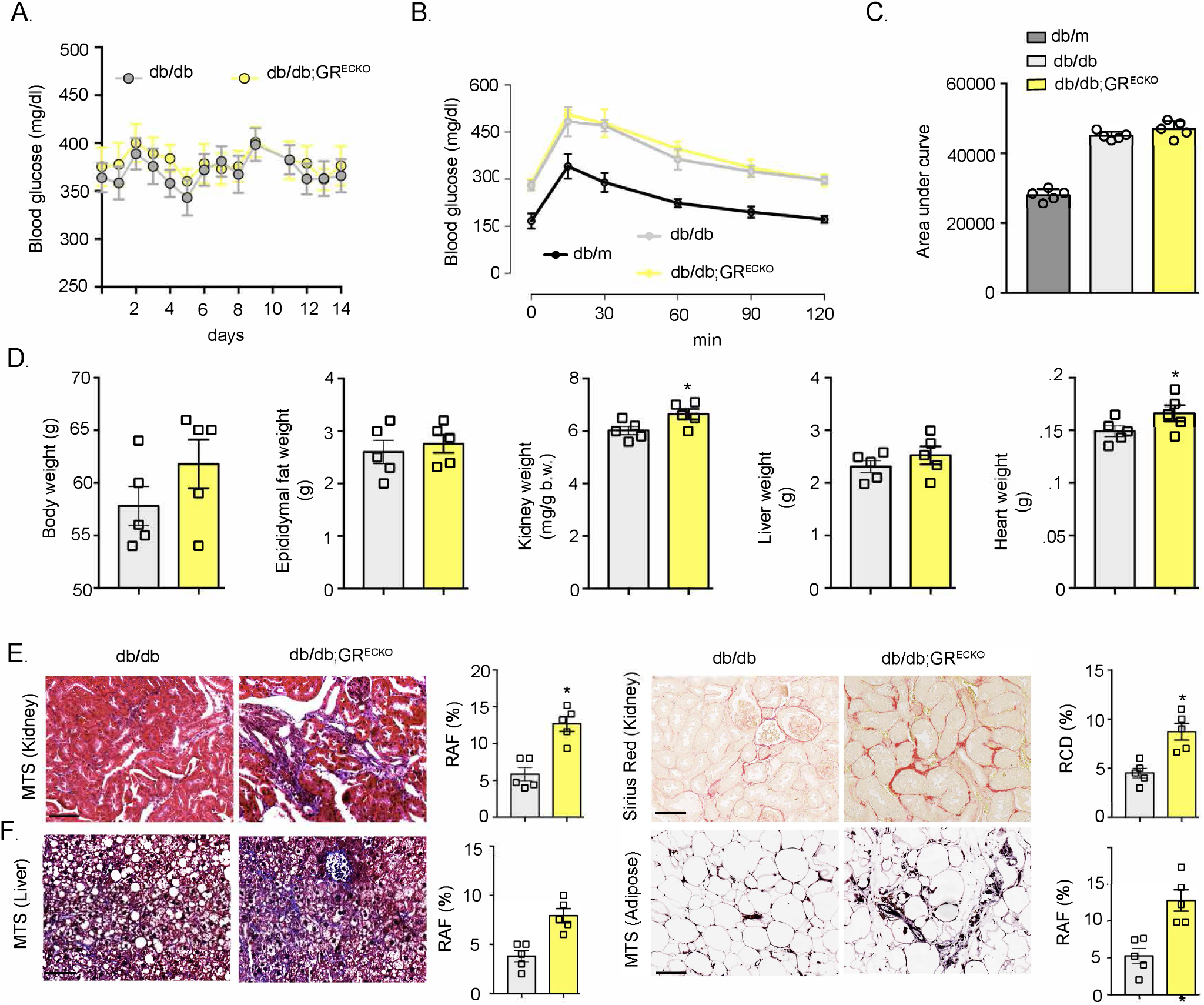
Loss of endothelial cell GR causes organ fibrosis in *db/db* mice. **(A)** Postprandial blood glucose measurements after 12-hour fasts over a 2-week period in db/db controls and endothelial cell-specific knock GR out in db/db mice (*db/db;GR^ECKO^*). n=5/group. **(B)** Glucose tolerance test (GTT) in control db/db mice and in *db/db;GR^ECKO^* mice. Heterozygous (non-diabetic) +/db (*db/m*) mice were included as a control. n=5/group. **(C)** AUC in control *db/db, db/m* and *db/db;GR^ECKO^* mice. n=5/group. **(D)** Physiological parameters including body weight, epididymal fat weight, kidney weight/body weight, liver weight and heart weight in control db/db and db/db;GR^ECKO^ mice. n=5/group. **(E)** MTS and Sirius red staining analysis in the kidneys of control db/db and db/db;GR^ECKO^ mice. Representative pictures at 30x magnification are shown. n=5/group. Relative area of fibrosis (RAF) and relative collagen deposition (RCD) were quantified using ImageJ. **(F)** MTS and Sirius red staining analysis in heart sections of control *db/db* and *db/db;GR^ECKO^* mice. Representative pictures at 20x are shown. n=5/group. **(G)** MTS in liver and **(H)** adipose tissue from control *db/db* and *db/db;GR^ECKO^* mice. Representative pictures at 20x are shown. n=5/group. Scale bar: 50μm. Data represent mean ± SEM. *P ≤ 0.05.

### Wnt inhibition improves organ fibrosis in diabetes

Next, we tested the effect of Wnt inhibition on glycemic control in db/db mice using a small molecule Wnt inhibitor, LGK974 (Wnti). Post-prandial glucose measurements after a 12-hour fast in *db/db* mice that were either untreated or treated with Wnti for 8 weeks revealed a trend toward improved glycemic control in treated mice (Figure 2A). A formal oral glucose tolerance test did show a small but significant improvement in glucose control in the Wnti-treated mice which was most pronounced at the later time points (Figure 2B,C). Additional phenotyping showed no differences between these 2 groups of mice (Figure 2D). Histologic examination of organs from untreated and Wnti-treated *db/db* mice showed a dramatic improvement in organ fibrosis in the Wnti-treated group (Figure 2E-F).

**Figure 2:**
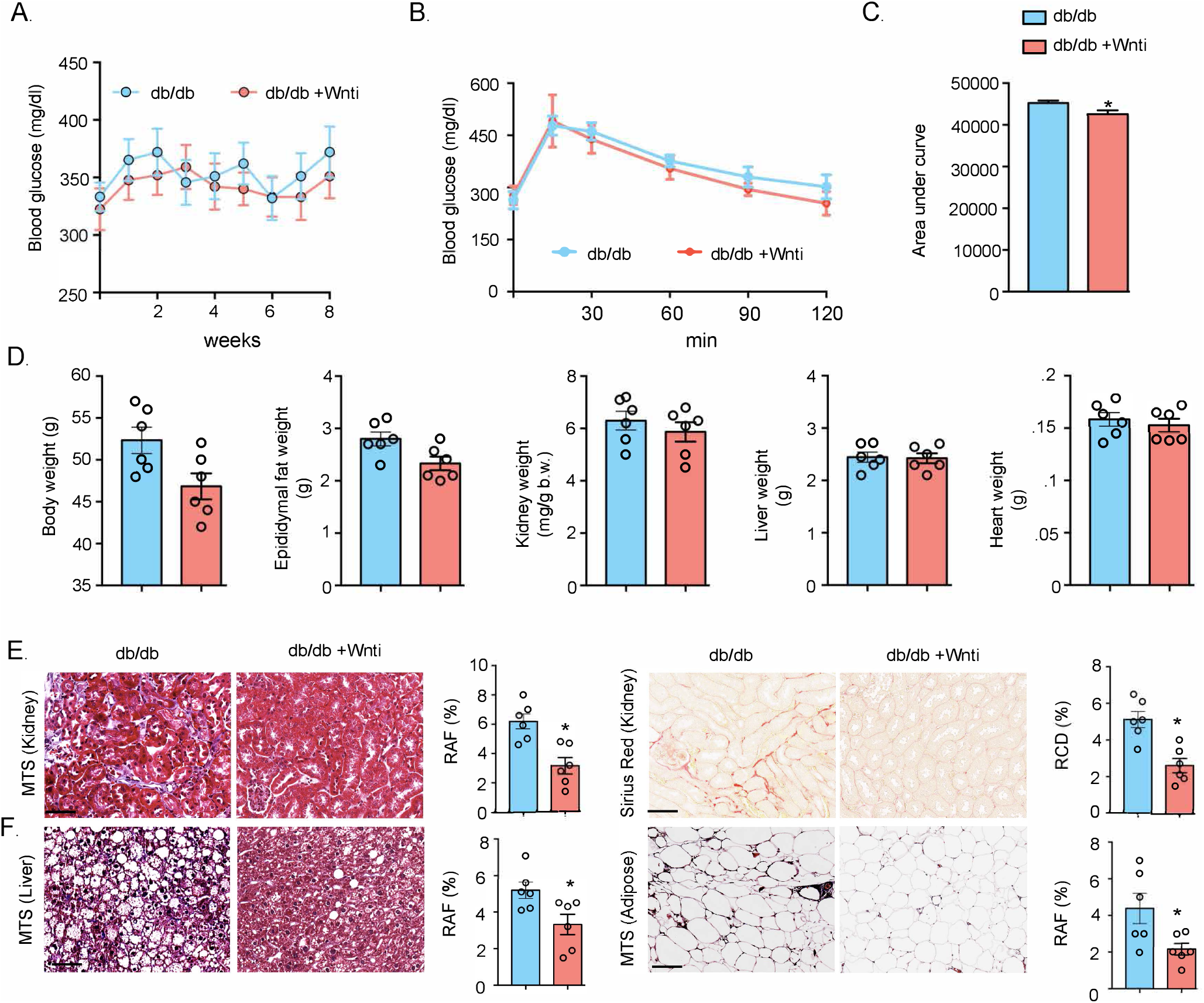
Pharmacological inhibition of Wnt signaling abolishes the fibrogenic phenotype in *db/db* mice. **(A)** Postprandial blood glucose measurements in the Wnt inhibitor treated *db/db* mice and untreated control *db/db* mice. LGK-974 (Wnti) was administered at 5 mg/kg body weight dorally by gavage. **(B)** Glucose tolerance test (GTT) analysis in the Wnti-treated *db/db* mice and untreated control *db/db* mice. n=6/group. **(C)** AUC for Wnti-treated *db/db* mice and untreated control *db/db* mice. n=6/group. **(D)** Physiological parameters such as body weight, epididymal fat, kidney weight/body weight, liver weight and heart weight in Wnti-treated *db/db* mice and untreated control *db/db* mice. n=6/group. **(E)** MTS and Sirius red staining in the kidneys of Wnti-treated *db/db* mice and untreated control *db/db* mice. Representative pictures at 30x are shown. n=6/group. Relative area of fibrosis (RAF) and relative collagen deposition (RCD) were quantified using ImageJ. **(F)** MTS staining in liver and adipose sections from Wnti-treated *db/db* mice and untreated control *db/db* mice. Representative pictures at 20x are shown. n=6/group. Scale bar: 50μm. Data represent the mean ± SEM. *P ≤ 0.05.

### Wnt inhibition is as effective as metformin in preventing organ fibrosis in HFD-fed mice

Prolonged HFD feeding leads to obesity, insulin resistance and eventually diabetes. Based on our previous work demonstrating a striking benefit of Wnt inhibition on renal fibrosis in streptozotocin-induced diabetes (5), we assessed whether blockade of Wnt signaling was able to temper fibrosis in HFD-fed control (*GR^fl/fl^, Cre-*) and *GR^ECKO^* mice. Metformin, an oral antihyperglycemic agent which improves insulin sensitivity and fibrosis in diabetes, was used as a positive control. Mice that had been HFD-fed for 20 weeks were either untreated, treated with Wnti for 8 weeks or treated with metformin for 8 weeks and then subjected to an oral glucose tolerance test. Interestingly, Wnt inhibition was able to partially blunt the hyperglycemic response, but not to the same extent as metformin (Figure 3A). A similar result was achieved when HFD-fed *GR^ECKO^* mice were exposed to the same treatments (Figure 3B). HFD-fed *GR^ECKO^* mice treated with either Wnti or metformin demonstrated lower body weight and decreased fat mass compared to similarly treated controls but no differences in organ weights were observed (Figure 3C). Histology from the kidney, heart, liver and adipose tissue from mice in each of the six treatment conditions was examined. Metformin, but not Wnti, was able to significantly improve fibrosis in all organs studied in the control animals, while in the *GR^ECKO^* mice both Wnti and metformin were able to significantly improve organ fibrosis (Figure 3D); the magnitude of the effect of metformin and Wnti in *GR^ECKO^* mice was similar.

**Figure 3:**
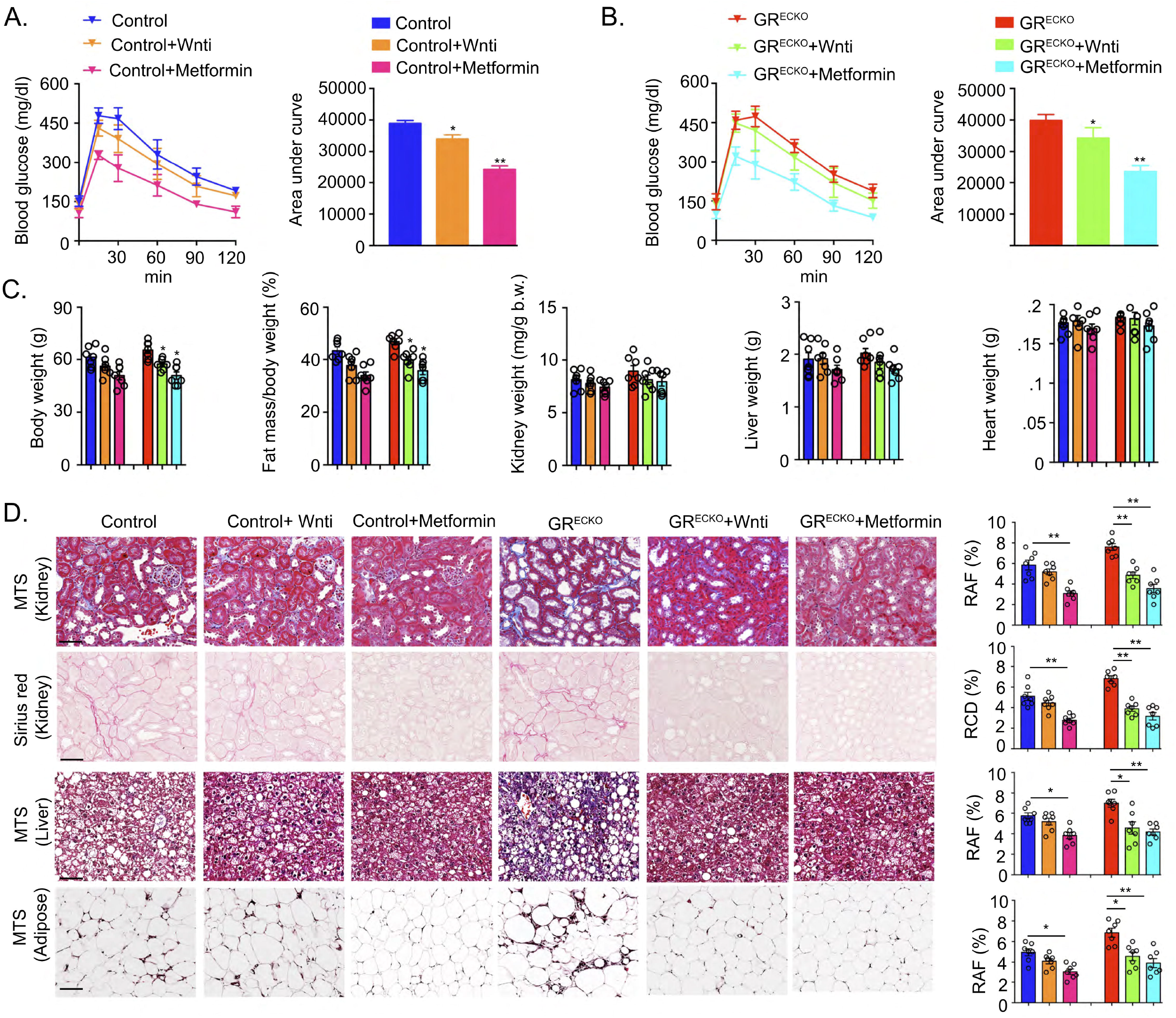
Effect of Wnti or metformin treatment in HFD-induced organ fibrosis. **(A)** GTT and AUC in HFD-fed control mice either untreated, treated with Wnti, or treated with metformin. n=7/group. **(B)** GTT and AUC in HFD-fed *GR^ECKO^* mice either untreated, treated with Wnti, or treated with metformin. n=7/group. Wnti was administered at 5 mg/kg body weight and metformin was administered at 100 mg/kg body weight. **(C)** Physiological parameters such as body weight, fat mass/body weight, kidney weight/body weight, liver weight and heart weight in the HFD-fed control or *GR^ECKO^* mice either untreated, treated with Wnti, or treated with metformin. n=7/group. **(D)** MTS and Sirius red staining from kidneys and heart sections. MTS staining from liver and adipose tissue is also shown. n=7/group. Relative area of fibrosis (RAF) and relative collagen deposition (RCD) were quantified using ImageJ. Scale bar: 50μm. Data represent mean ± SEM. *P ≤ 0.05 and **P ≤ 0.01.

### IL-6 is a key cytokine driving the fibrosis phenotype

In order to investigate further the mechanism of fibrosis in these animals, we isolated plasma from control and *GR^ECKO^* mice on both a standard chow diet and HFD and assessed the levels of key inflammatory cytokines including IL-1β, IL-6 and IFN-ý. HFD-fed animals of both genotypes demonstrated higher levels of IL-1β and IL-6, with HFD-fed *GR^ECKO^* mice demonstrating the highest levels of all (Figure 4A). There were no significant differences in IFN-ý levels. Next, plasma from control and *GR^ECKO^* mice, either untreated, treated with Wnti or treated with metformin, was tested. IL-1β and IL-6 levels were again highest in *GR^ECKO^* mice but could be suppressed by about 50% both with Wnti treatment and metformin treatment, comparable to the levels of similarly treated controls (Figure 4B). To examine this phenomenon in a cell culture system, HUVECs were treated with either control siRNA alone, GR siRNA alone or GR siRNA and an IL-6 neutralization antibody (N-IL-6). As shown in Figure 4C, GR siRNA-treated cells had the highest levels of IL-6 and highest gene expression of the associated fibrogenic markers alpha-SMA and FSP-1 as well as the Wnt-dependent gene *axin2;* the expression of all of these was significantly suppressed with the administration of N-IL-6.

**Figure 4:**
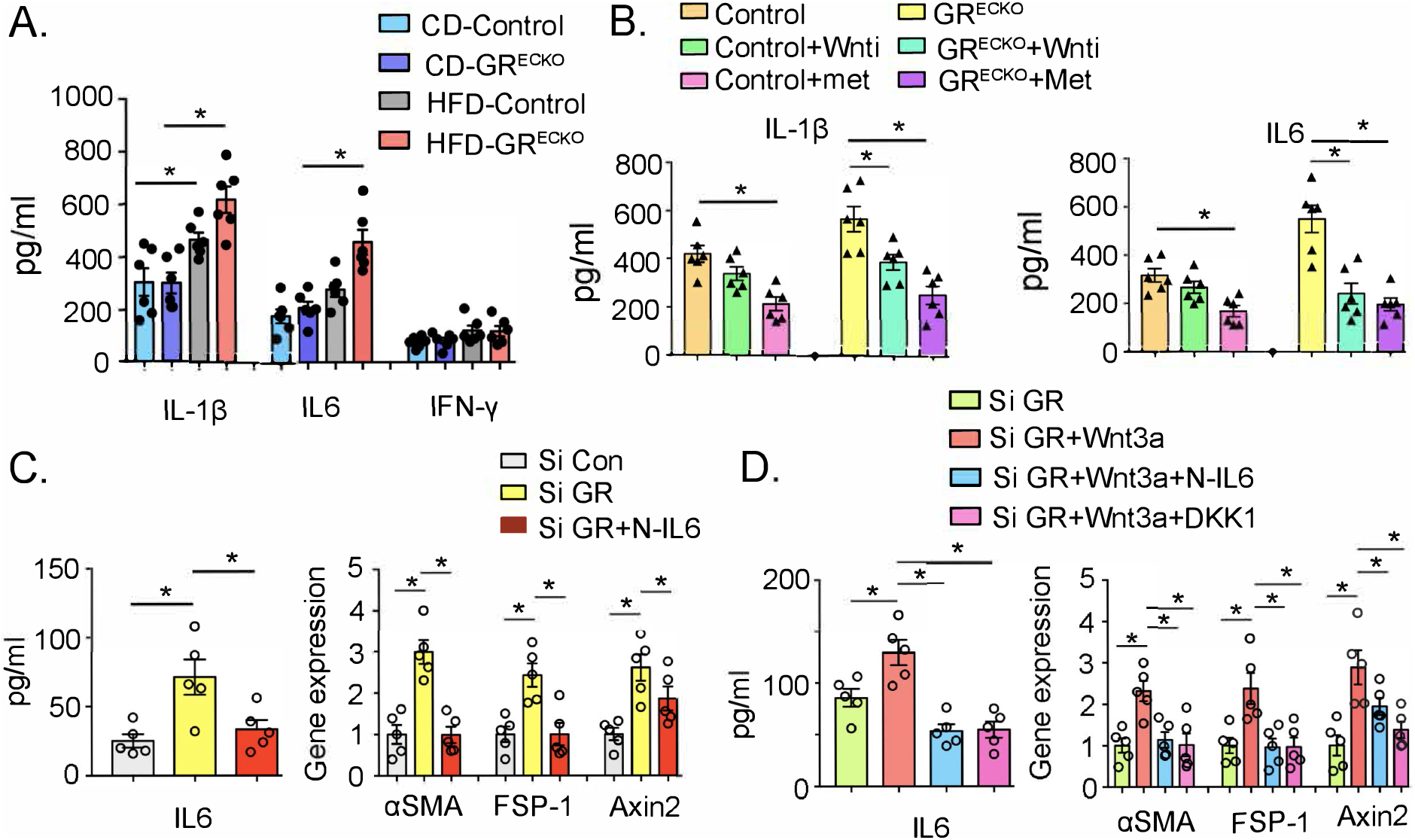
Both GR loss and Wnt activation are associated with pro-inflammatory and pro-fibrotic signaling pathways. **(A)** IL-1β, IL-6 and IFN-γ were measured in plasma from control diet (CD)- and HFD-fed control and *GR^ECKO^* mice using Luminex cytokine array analysis. n=6/group. **(B)** IL-1β and IL-6 levels from plasma from HFD-fed control mice treated with Wnti, metformin, or untreated and HFD-fed *GR^ECKO^* mice treated with Wnti, metformin, or untreated are shown. n=6/group. **(C)** IL-6 levels in the media of cultured HUVECs treated with either control siRNA (si Con), GR siRNA (si GR) or GR siRNA with neutralization antibody of IL-6 (si GR+N-IL6) are shown. Relative expression of the indicated genes was normalized to 18S. **(D)** IL-6 levels in conditioned media from GR siRNA-treated HUVECs treated with recombinant Wnt3a alone or in combination with either N-IL-6- or DKK-1-are shown. Relative expression of the indicated genes was normalized to 18S. Data are mean ± SEM. *p < 0.05.

Finally, to determine if the IL-6-mediated fibrotic effects were mechanistically tied to Wnt signaling, the GR siRNA-treated cells were either left untreated, or concurrently treated with the canonical Wnt ligand Wnt3a (alone), Wnt3a and N-IL-6 or Wnt3a and DKK-1, an inhibitor of canonical Wnt. Administration of Wnt3a alone significantly increased IL-6 levels which could be similarly suppressed with either N-IL-6 or DKK-1. The mRNA expression of fibrogenic markers and *axin2* was significantly increased by Wnt3a and could be similarly suppressed by either N-IL-6 or DKK-1, suggesting that IL-6 expression is directly linked to Wnt signaling.

## Discussion

The major finding of this work is that endothelial GR-mediated upregulation of Wnt signaling and concurrent hyperinflammation work synergistically to exacerbate organ fibrosis in a model of diabetes. This phenomenon is analogous to the ‘two-hit’ hypothesis that is observed in other conditions with renal manifestations such as autosomal dominant polycystic kidney disease (ADPKD) (11), tuberous sclerosis (12) and von Hippel-Lindau disease (13). Recent work also demonstrates that non-alcoholic fatty liver disease (NAFLD), the risk of which is heightened by the presence of type 2 diabetes, is characterized by a ‘two-hit’ or even a ‘multi-hit’ hypothesis which also includes a hyperinflammatory state and dysregulated Wnt signaling in addition to triglyceride accumulation in hepatocytes (14).

Metformin is an old and inexpensive drug that remains first-line therapy for type 2 diabetes mellitus. However, recognition of its pleiotropic effects has suggested that it may be beneficial in a wide variety of chronic diseases. For example, data from both animals and humans support the notion that metformin can modulate inflammation (15). Metformin is able to shift the balance of macrophages in favor of the M2 subtype which decreases the release of cytokines such as TNF-alpha, IL-beta and IL-6, and heightens the anti-inflammatory response (16). Metformin was recently shown to suppress high-fat diet-induced upregulated Wnt signaling in a mouse model of age-related leaky gut and inflammation (17).

With regard to use in kidney disease, metformin has been shown to be associated with suppression of oxidative stress and inflammation in a rat model of diabetic nephropathy (18). In this study, the authors demonstrated these improvements were specifically mediated by the inhibition of renal artery angiotensin II receptor type I/endothelin-1 axis (18), suggesting that much of the damage in diabetic nephropathy in this model was mediated via vascular effects. A beneficial effect of metformin on the progression of ADPKD was recently shown by Pastor-Soler and colleagues, particularly in the reduction of key inflammatory markers and cell proliferation markers (19). One of the major limitations to broader use of metformin as a therapy for fibrosis is that the doses required often lead to clinically-significant hypoglycemia. Indeed, in our study we did observe a 30-40% decrease in peak glucose concentration after glucose tolerance testing in HFD-fed, metformin-treated mice compared to controls. One potential solution to this problem is the development of a targeted drug delivery systems which involves the synthesis of nanoparticles comprised of grafted chitosan which acts as a metformin carrier and can be specifically delivered to renal tubular cells by megalin-mediated endocytosis (20).

A limitation of this study is that only male *db/db* mice were studied. There is literature to support a sex-specific effect of metformin (21). For example, metformin is generally associated with some degree of cardio-protection in women but not necessarily in men (22, 23). Conversely, metformin has been shown to prevent and reverse neuropathic pain only in male mice (24). It is unclear if the effects of metformin in female *db/db*; *GR^ECKO^* would have been the same as those observed in the male mice.

In conclusion, the *db/db*; *GR^ECKO^* mouse model is a unique model in which the contributions of diabetes and endogenous cortisol signaling can be studied simultaneously. There is much we can learn with regard to how the tonic anti-inflammatory effects of endothelial cortisol act as a ‘brake’ on organ fibrosis and unchecked vascular inflammation.

## Author Contributions

S.P.S. and J.E.G. conceived and designed research. S.P.S. performed experiments. S.P.S. and J.E.G. analyzed data and prepared figures. J.E.G. drafted the manuscript.

## References

1. Reidy K, Kang HM, Hostetter T, Susztak K. Molecular mechanisms of diabetic kidney disease. J Clin Invest. 2014;124(6):2333–40. Epub 2014/06/04. doi: 10.1172/JCI72271. PubMed PMID: 24892707; PMCID: PMC4089448.

2. Badal SS, Danesh FR. New insights into molecular mechanisms of diabetic kidney disease. Am J Kidney Dis. 2014;63(2 Suppl 2):S63–83. Epub 2014/01/28. doi: 10.1053/j.ajkd.2013.10.047. PubMed PMID: 24461730; PMCID: PMC3932114.

3. Tesch GH, Lim AK. Recent insights into diabetic renal injury from the db/db mouse model of type 2 diabetic nephropathy. Am J Physiol Renal Physiol. 2011;300(2):F301–10. Epub 2010/12/15. doi: 10.1152/ajprenal.00607.2010. PubMed PMID: 21147843.

4. Cruz TB, Carvalho FA, Matafome PN, Soares RA, Santos NC, Travasso RD, Oliveira MJ. Mice with Type 2 Diabetes Present Significant Alterations in Their Tissue Biomechanical Properties and Histological Features. Biomedicines. 2021;10(1). Epub 2022/01/22. doi: 10.3390/biomedicines10010057. PubMed PMID: 35052737; PMCID: PMC8773308.

5. Srivastava SP, Zhou H, Setia O, Liu B, Kanasaki K, Koya D, Dardik A, Fernandez-Hernando C, Goodwin J. Loss of endothelial glucocorticoid receptor accelerates diabetic nephropathy. Nat Commun. 2021;12(1):2368. Epub 2021/04/24. doi: 10.1038/s41467-021-22617-y. PubMed PMID: 33888696; PMCID: PMC8062600.

6. Zhou H, Mehta S, Srivastava SP, Grabinska K, Zhang X, Wong C, Hedayat A, Perrotta P, Fernandez-Hernando C, Sessa WC, Goodwin JE. Endothelial cell-glucocorticoid receptor interactions and regulation of Wnt signaling. JCI Insight. 2020;5(3). Epub 2020/02/14. doi: 10.1172/jci.insight.131384. PubMed PMID: 32051336; PMCID: PMC7098785.

7. Schunk SJ, Floege J, Fliser D, Speer T. WNT-beta-catenin signalling - a versatile player in kidney injury and repair. Nat Rev Nephrol. 2021;17(3):172–84. Epub 2020/09/30. doi: 10.1038/s41581-020-00343-w. PubMed PMID: 32989282.

8. Goodwin JE, Feng Y, Velazquez H, Sessa WC. Endothelial glucocorticoid receptor is required for protection against sepsis. Proc Natl Acad Sci U S A. 2013;110(1):306–11. Epub 2012/12/19. doi: 10.1073/pnas.1210200110. PubMed PMID: 23248291; PMCID: PMC3538225.

9. Goodwin JE, Zhang X, Rotllan N, Feng Y, Zhou H, Fernandez-Hernando C, Yu J, Sessa WC. Endothelial glucocorticoid receptor suppresses atherogenesis--brief report. Arterioscler Thromb Vasc Biol. 2015;35(4):779–82. Epub 2015/03/27. doi: 10.1161/ATVBAHA.114.304525. PubMed PMID: 25810297; PMCID: PMC4375730.

10. Zhang LS, Lum L. Delivery of the Porcupine Inhibitor WNT974 in Mice. Methods Mol Biol. 2016;1481:111–7. doi: 10.1007/978-1-4939-6393-5_12. PubMed PMID: 27590157; PMCID: PMC5024565.

11. Nauli SM, Rossetti S, Kolb RJ, Alenghat FJ, Consugar MB, Harris PC, Ingber DE, Loghman-Adham M, Zhou J. Loss of polycystin-1 in human cyst-lining epithelia leads to ciliary dysfunction. J Am Soc Nephrol. 2006;17(4):1015–25. Epub 2006/03/28. doi: 10.1681/ASN.2005080830. PubMed PMID: 16565258.

12. Wilson C, Bonnet C, Guy C, Idziaszczyk S, Colley J, Humphreys V, Maynard J, Sampson JR, Cheadle JP. Tsc1 haploinsufficiency without mammalian target of rapamycin activation is sufficient for renal cyst formation in Tsc1+/-mice. Cancer Res. 2006;66(16):7934–8. Epub 2006/08/17. doi: 10.1158/0008-5472.CAN-06-1740. PubMed PMID: 16912167.

13. van der Horst-Schrivers ANA, Sluiter WJ, Kruizinga RC, van Leeuwaarde RS, Giles R, Olderode-Berends MJW, Links TP. The incidence of consecutive manifestations in Von Hippel-Lindau disease. Fam Cancer. 2019;18(3):369–76. Epub 2019/05/16. doi: 10.1007/s10689-019-00131-x. PubMed PMID: 31087189; PMCID: PMC6560011.

14. Pan J, Zhou W, Xu R, Xing L, Ji G, Dang Y. Natural PPARs agonists for the treatment of nonalcoholic fatty liver disease. Biomed Pharmacother. 2022;151:113127. Epub 2022/05/23. doi: 10.1016/j.biopha.2022.113127. PubMed PMID: 35598367.

15. Bharath LP, Agrawal M, McCambridge G, Nicholas DA, Hasturk H, Liu J, Jiang K, Liu R, Guo Z, Deeney J, Apovian CM, Snyder-Cappione J, Hawk GS, Fleeman RM, Pihl RMF, Thompson K, Belkina AC, Cui L, Proctor EA, Kern PA, Nikolajczyk BS. Metformin Enhances Autophagy and Normalizes Mitochondrial Function to Alleviate Aging-Associated Inflammation. Cell Metab. 2020;32(1):44–55 e6. Epub 2020/05/14. doi: 10.1016/j.cmet.2020.04.015. PubMed PMID: 32402267; PMCID: PMC7217133.

16. Jing Y, Wu F, Li D, Yang L, Li Q, Li R. Metformin improves obesity-associated inflammation by altering macrophages polarization. Mol Cell Endocrinol. 2018;461:256–64. Epub 2017/09/25. doi: 10.1016/j.mce.2017.09.025. PubMed PMID: 28935544.

17. Ahmadi S, Razazan A, Nagpal R, Jain S, Wang B, Mishra SP, Wang S, Justice J, Ding J, McClain DA, Kritchevsky SB, Kitzman D, Yadav H. Metformin Reduces Aging-Related Leaky Gut and Improves Cognitive Function by Beneficially Modulating Gut Microbiome/Goblet Cell/Mucin Axis. J Gerontol A Biol Sci Med Sci. 2020;75(7):e9–e21. Epub 2020/03/05. doi: 10.1093/gerona/glaa056. PubMed PMID: 32129462; PMCID: PMC7302182.

18. Dawood AF, Maarouf A, Alzamil NM, Momenah MA, Shati AA, Bayoumy NM, Kamar SS, Haidara MA, ShamsEldeen AM, Yassin HZ, Hewett PW, Al-Ani B. Metformin Is Associated with the Inhibition of Renal Artery AT1R/ET-1/iNOS Axis in a Rat Model of Diabetic Nephropathy with Suppression of Inflammation and Oxidative Stress and Kidney Injury. Biomedicines. 2022;10(7). Epub 2022/07/28. doi: 10.3390/biomedicines10071644. PubMed PMID: 35884947; PMCID: PMC9313150.

19. Pastor-Soler NM, Li H, Pham J, Rivera D, Ho PY, Mancino V, Saitta B, Hallows KR. Metformin improves relevant disease parameters in an autosomal dominant polycystic kidney disease mouse model. Am J Physiol Renal Physiol. 2022;322(1):F27–F41. Epub 2021/11/23. doi: 10.1152/ajprenal.00298.2021. PubMed PMID: 34806449.

20. Sun H, Shi K, Zuo B, Zhang X, Liu Y, Sun D, Wang F. Kidney-Targeted Drug Delivery System Based on Metformin-Grafted Chitosan for Renal Fibrosis Therapy. Mol Pharm. 2022;19(9):3075–84. Epub 2022/08/09. doi: 10.1021/acs.molpharmaceut.1c00827. PubMed PMID: 35938707.

21. Ilias I, Rizzo M, Zabuliene L. Metformin: Sex/Gender Differences in Its Uses and Effects-Narrative Review. Medicina (Kaunas). 2022;58(3). Epub 2022/03/27. doi: 10.3390/medicina58030430. PubMed PMID: 35334606; PMCID: PMC8952223.

22. Lyons MR, Peterson LR, McGill JB, Herrero P, Coggan AR, Saeed IM, Recklein C, Schechtman KB, Gropler RJ. Impact of sex on the heart’s metabolic and functional responses to diabetic therapies. Am J Physiol Heart Circ Physiol. 2013;305(11):H1584–91. Epub 2013/09/18. doi: 10.1152/ajpheart.00420.2013. PubMed PMID: 24043256; PMCID: PMC3882467.

23. Raparelli V, Elharram M, Moura CS, Abrahamowicz M, Bernatsky S, Behlouli H, Pilote L. Sex Differences in Cardiovascular Effectiveness of Newer Glucose-Lowering Drugs Added to Metformin in Type 2 Diabetes Mellitus. J Am Heart Assoc. 2020;9(1):e012940. Epub 2020/01/07. doi: 10.1161/JAHA.119.012940. PubMed PMID: 31902326; PMCID: PMC6988160.

24. Inyang KE, Szabo-Pardi T, Wentworth E, McDougal TA, Dussor G, Burton MD, Price TJ. The antidiabetic drug metformin prevents and reverses neuropathic pain and spinal cord microglial activation in male but not female mice. Pharmacol Res. 2019;139:1–16. Epub 2018/11/06. doi: 10.1016/j.phrs.2018.10.027. PubMed PMID: 30391353; PMCID: PMC6447087.

